# Verkko2: Integrating proximity ligation data with long-read De Bruijn graphs for efficient telomere-to-telomere genome assembly, phasing, and scaffolding

**DOI:** 10.1101/2024.12.20.629807

**Authors:** Dmitry Antipov, Mikko Rautiainen, Sergey Nurk, Brian P. Walenz, Steven J. Solar, Adam M. Phillippy, Sergey Koren

## Abstract

The Telomere-to-Telomere Consortium recently finished the first truly complete sequence of a human genome. To resolve the most complex repeats, this project relied on the semi-manual combination of long, accurate PacBio HiFi and ultra-long Oxford Nanopore sequencing reads. The Verkko assembler later automated this process, achieving complete assemblies for approximately half of the chromosomes in a diploid human genome. However, the first version of Verkko was computationally expensive and could not resolve all regions of a typical human genome. Here we present Verkko2, which implements a more efficient read correction algorithm, improves repeat resolution and gap closing, introduces proximity-ligation-based haplotype phasing and scaffolding, and adds support for multiple long-read data types. These enhancements allow Verkko to assemble all regions of a diploid human genome, including the short arms of the acrocentric chromosomes and both sex chromosomes. Together, these changes increase the number of telomere-to-telomere scaffolds by twofold, reduce runtime by fourfold, and improve assembly correctness. On a panel of 19 human genomes, Verkko2 assembles an average of 39 of 46 complete chromosomes as scaffolds, with 21 of these assembled as gapless contigs. Together, these improvements enable telomere-to-telomere comparative and pangenomics, at scale.

## Introduction

Advances in sequencing technologies have set a new benchmark for *de novo* genome assembly — the complete assembly of entire chromosomes from ‘telomere to telomere’ (T2T), defined here as being complete, gapless, and containing telomere on both ends. The current sequencing recipe for achieving T2T assemblies combines ‘long accurate reads’ (LA reads) of length *>* 10 *kb* and accuracy greater than 99.9% with ‘ultra-long reads’ (UL reads) of length *>* 100 *kb* and accuracy higher than 95%. Chromosome-scale haplotype separation is then achieved using either parental (Koren *et al*. 2018) or Hi-C sequencing (Cheng *et al*. 2021; Garg 2021) data, resulting in the complete assembly of both maternal and paternal haplotypes. Verkko (Rautiainen *et al*. 2023) previously demonstrated the power of this recipe, automatically assembling 20 of 46 diploid human chromosomes without gaps. While subsequent tools have offered improved performance (Cheng *et al*. 2024) and a reduced dependence on trios (Lorig-Roach *et al*. 2024), no assembler can yet assemble all chromosomes of a typical human genome. Genomic features that remain a challenge, even for ultra-long-read sequencing, include many functionally relevant regions of the genome such as recent segmental duplications, tandemly duplicated satellite repeats, and amplified gene arrays (e.g. rDNAs).

The short arms of the human acrocentric chromosomes, whose base-level structure was first revealed by the T2T-CHM13 reference genome (Nurk *et al*. 2022), are most challenging to assemble due to their long, tandemly repeated rDNA arrays and enrichment for segmental duplications. Translocations (Ohno *et al*. 1961) and ongoing recombination (Guarracino *et al*. 2023) within the short arms have maintained a high sequence similarity across all five acrocentric chromosomes (Nurk *et al*. 2022), and the rDNA arrays can stretch for megabases, exceeding the length of current ultra-long sequencing reads. Thus, correctly resolving the acrocentrics in a diploid genome requires differentiating between ten similar short arms and scaffolding across each rDNA array gap. Doing so is necessary for understanding variation within these important regions of the genome and better detecting abnormalities, such as Robertsonian chromosomes (de Lima *et al*. 2024).

The distal regions of the short arms are typically connected in the assembly graph (Figure 1), and standard Hi-C phasing algorithms that expect a diploid ‘bubble chain’ structure fail to separate them by haplotype (Kronenberg *et al*. 2021; Cheng *et al*. 2022). Without prior separation of chromosomes by haplotype, single-haplotype Hi-C scaffolders (Zhou *et al*. 2023; Ghurye *et al*. 2019; Burton *et al*. 2013) cannot be used to bridge the rDNA gaps. Furthermore, existing tools specifically designed for diploid Hi-C scaffolding (Zhang *et al*. 2019; Zeng *et al*. 2024) are also unsuitable for these complex and repetitive regions of the genome because they do not make use of the assembly graph nor consider multi-mapped reads. This problem is not unique to human, and other species such as sheep and cattle have many more acrocentric or telocentric chromosomes than human (23 and 29, respectively) that also share a high degree of sequence similarity on their ends (Kalbfleisch *et al*. 2024). Enabling the T2T assembly of more samples and species requires a generalized solution for dealing with such inter-chromosomal sequence homology.

**Fig. 1:**
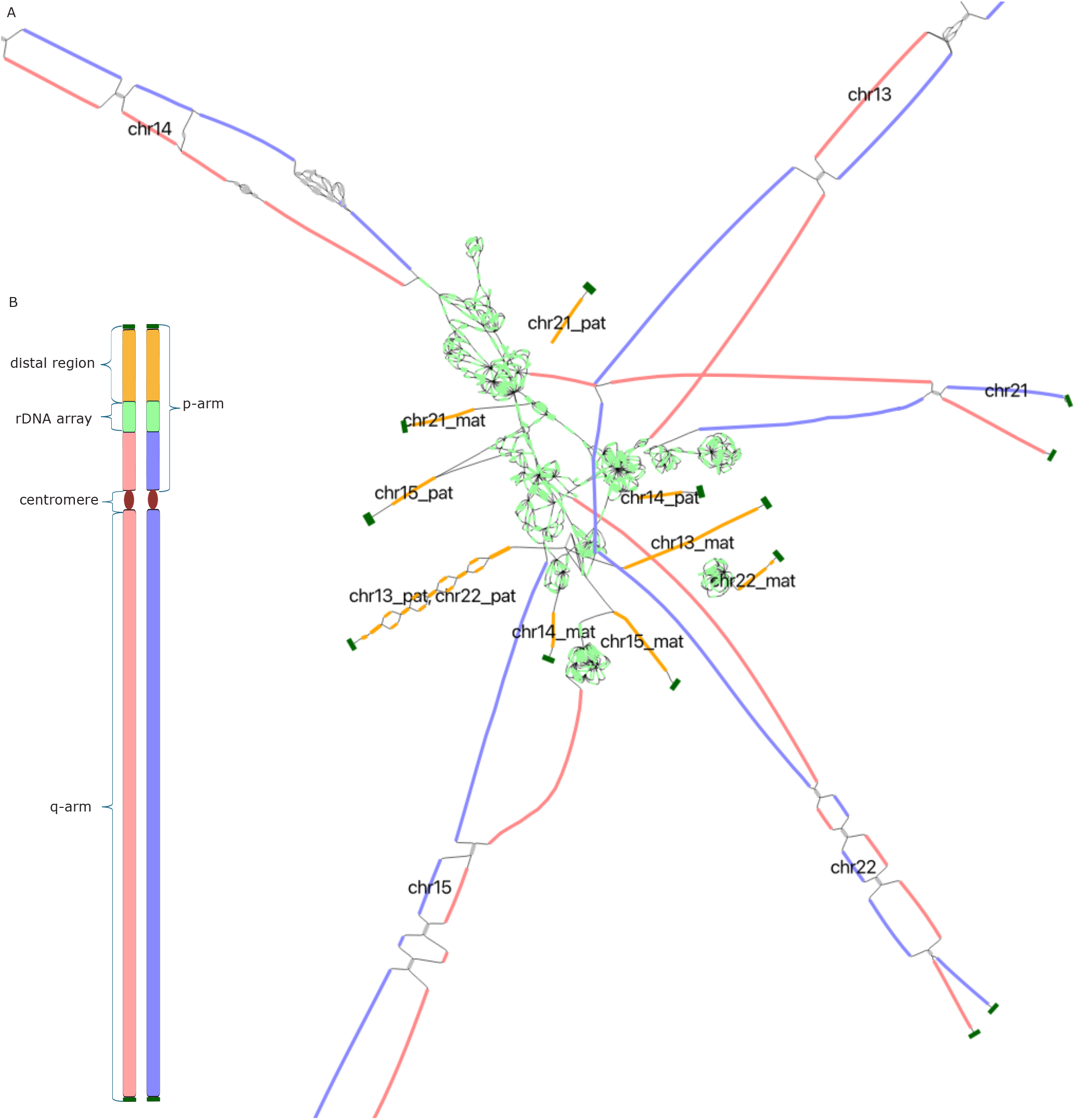
(A) Assembly graph tangle resulting from the rDNA arrays of the HG002 human genome. Bandage (Wick *et al*. 2015) visualization of the assembly graph for the ten HG002 acrocentric chromosomes (diploid chromosomes 13, 14, 15, 21, 22). Maternal and paternal haplotype-assigned nodes are shown in red and blue, respectively, the rDNA repeats in light green, the distal satellite regions in orange, and the telomeres in dark green. Each distal satellite is labeled according to the HG002 v1.0.1 reference assembly, with the exception of Chromosome 13 and Chromosome 22 paternal which are too similar to be separated (Potapova *et al*. 2024a). (B) Linear schematic of a human acrocentric chromosome with colors matching panel A. The paternal and maternal haplotype-assigned sequences comprise the entire q-arm and the proximal component of the p-arm. In the contrast to other regions of the genome, the short arms of the acrocentric chromosomes do not split into a typical diploid arrangement and are difficult to phase correctly. Such complex structures violate the assumptions made by typical diploid phasing and scaffolding tools.

We have developed a new version of Verkko to address these issues. In addition to a dramatically accelerated read-correction step, Verkko2 natively supports Hi-C data to enable the assembly of genomes where trio information is not available. Verkko2 also introduces a new scaffolding module that is robust to the complex graph structures created by acrocentric chromosomes. Rather than scaffolding individual segments of assembly graph (Ghurye *et al*. 2019), Verkko2 scaffolds haplotype paths after Hi-C or trio-based phasing. This scaffolding step considers both uniquely and multi-mapped reads, making minimal assumptions regarding the graph structure, allowing for scaffolding in even the most repetitive regions of the genome. Lastly, Verkko2 tracks every read through the assembly process and provides a BAM file for each assembled contig, detailing the reads used, their location in the assembly, and their base-level alignment to the final sequence. This removes the need for read re-mapping during downstream consensus polishing and validation processes (Mc Cartney *et al*. 2022; Liao *et al*. 2023; Mastoras *et al*. 2024a). Below we describe these changes in detail and evaluate Verkko2 on 19 human and two non-human samples, demonstrating consistent improvements in speed, continuity, and accuracy.

## Methods

The original Verkko pipeline consisted of six main steps, with module dependencies given in parentheses:

- Homopolymer compression and correction of LA reads (HiCanu, Nurk *et al*. (2020))
- LA multiplex De Bruijn graph construction (MBG, Rautiainen and Marschall (2021))
- Alignment of UL reads to the LA graph (GraphAligner, Rautiainen and Marschall (2020))
- ULA multiplex De Bruijn graph construction
- Haplotype path extension using haplotype-specific markers
- Haplotype path consensus generation from raw LA and UL reads (Canu, Koren *et al*. (2017))

Our improvements to Verkko are detailed below in order of their pipeline execution. Supplementary Figure S1 includes a graphical representation of original pipeline, adapted from (Rautiainen *et al*. 2023), further highlighting which stages of the Verkko pipeline were modified. All evaluations and comparisons described here are based on Verkko branch v1 fxd (Verkko1) and release v2.2.1 (Verkko2).

### Long, accurate read correction

Verkko previously re-used HiCanu’s read correction module to correct the LA reads (Nurk *et al*. 2020). This step first computed all-vs-all read overlaps followed by error correction based on the resulting alignments. The initial all-vs-all computation was a performance bottleneck and used as much as 79% of total runtime for the HG002 human genome assembly (Rautiainen *et al*. 2023). Verkko2 introduces a more efficient overlapper to dramatically reduce runtime.

The algorithm used by the new overlapper is based on long-*k* minimizer matching (Roberts *et al*. 2004; Jain *et al*. 2017). First, the reads are homopolymer and microsatellite compressed (Rautiainen *et al*. 2023) to mask systematic sequencing errors. Then, minimizers are extracted from each read using a 64-bit rolling hash and minimizer parameters of *k* = 425 and *w* = 19. The sequences of the minimizers are ignored and only the hash value and start and end positions within the read are stored. A maximum copy count (default 1, 000) is used to filter out repetitive minimizers. Matches per read are found based on the hash values and positions. The same minimizer occurring in two different reads implies a match between the reads, with the start positions of the minimizers specifying the match diagonal. The minimizer matches are sorted by diagonal and clustered such that any diagonal difference of at least 500 b between adjacent matches separates them into different clusters. Each cluster is then considered one overlap, with the start and end positions of the overlap taken from the minimum and maximum start and end positions of the minimizers in the cluster. A minimum overlap length threshold (default 1, 000 b) is used to remove spurious overlaps. Although the use of a 64-bit hash can introduce spurious matches, it is unlikely to cause spurious overlaps since that would require two or more hash collisions on similar match diagonals.

The implementation of the overlapper uses a disk index to reduce memory use. To build the index, the overlapper uses several temporary files: one *temporary k-mer file* and multiple *temporary hash files* where the number of temporary hash files *f* is given as a parameter (default *f* = 16). First all reads are iterated, minimizers are extracted, and their hashes and positions are stored in the temporary *k*-mer and hash files, respectively. The hashes are divided into the temporary hash files based on their values, with *f* temporary files placing the hash *h* into the *h* mod *f*’th file. Each temporary hash file is processed one at a time, where the occurrences of each hash value are counted and any hashes appearing only once are discarded. Then, all hashes appearing at least twice are assigned a unique incremental ID, with the first hash in the first file assigned *ID* = 0. The assigned hash IDs are stored in memory as a hash table from *k*-mer hashes to IDs. Finally, the temporary *k*-mer file is iterated to build the index file, where the hashes in the temporary *k*-mer file are replaced with their IDs and the tuple of (ID, read, start, end) is stored in the index file. Due to the read-by-read iteration when reading the *k*-mers, the *k*-mers in the index file will have the *k*-mers of a single read in a contiguous block.

Given *n* reads with *l* base pairs each, minimizer parameters *k* and *w*, number of temporary hash files *f*, and number of distinct hashes with copy count at least two *h*, the index construction takes worst case *O*(*nl*) time, and 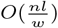 temporary disk space, disk writes and reads, and produces an index file of size 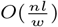. The expected index construction memory use is 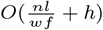 where the first term is the memory use for counting hash multiplicity and assigning IDs to the hashes, and the second term is the memory used for keeping the mapping from hashes to IDs. The value of *h* is typically small and close to genome size divided by *w* since it is composed of *k*-mers which appear at least twice in the reads, excluding most spurious *k*-mers.

To find the overlaps between the reads, the index file is iterated in multiple batches. Each batch inputs parameters *batch count c* and *batch index i*. The parameters are used to split the reads into *c* equally large ranges, with the *i*’th range indexed and matched in one batch. To find the overlaps, the index file is iterated and the *k*-mers of the reads in the indexed range *i* are stored in memory. Then, the index file is iterated again and whenever a read in range *x* ≤ *i* is encountered, all overlaps against the in-memory reads are computed and output. Since the *k*-mers of each read are in a contiguous block, the *k*-mers of the matched reads do not need to be stored in memory except for the single read currently being processed, and so the memory use is the index size divided by *c*. The batch count parameter provides a time-memory trade-off, with more batches requiring less memory but more passes through the index file. In total *c* batches are required to find all overlaps. Once all-vs-all overlaps are computed, Verkko re-uses the previous code to compute base-level alignments and extend each overlap beyond the *k*-mer matches.

### Repeat identification

As shown in (Rautiainen *et al*. 2023), Verkko performs well for the majority of the genome. However, some regions were prone to issues in Verkko1. The first improvement was to tolerate coverage fluctuations in the input LA sequencing reads. Both HiFi and ONT Duplex sequences, which are the most commonly used LA datatypes, have biases which lead to reduced coverage in certain contexts (Nurk *et al*. 2020; Koren *et al*. 2024). A critical step for repeat resolution is the initial identification of repeat and unique nodes by coverage to create source and sink nodes for subsequent ONT-based resolution. Rather than relying on absolute average coverage, Verkko1 attempted to correct for uneven coverage based on the surrounding graph neighborhood compared to the global graph average. We updated this criteria to compare against the local connected component average to better resolve regions with systematically lower or higher coverage.

### Telomere assembly

Another region Verkko1 struggled with was the telomere or the terminal node in a connected component in the graph. In many cases, assemblies would extend up to the telomere but miss the terminal node. Two issues were primarily responsible for this loss of sequence. First, tip clipping could remove multiple tip nodes during simplification, leaving no telomere sequence in the graph. Instead, we now ensure at least one tip node is retained, preferring to keep the highest-coverage or, when all nodes are equal coverage, the longest node. Second, the identification of unique nodes did not flag the last node in a component as unique since these nodes frequently have lower than expected coverage or are short. This prevented subsequent simplification from reaching the telomere, leaving an ambiguous branch in the graph. Verkko2 relaxes the uniqueness criteria for dead-end nodes to ensure that these branches are resolved.

### Gap closing

While investigating gaps in Verkko1 assemblies, we found that GraphAligner could systematically misalign reads to the wrong haplotype. In cases where one haplotype had missing sequence, due to a lack of LA coverage. The correct alignment requires splitting a read into at least two pieces and leaving bases unaligned. As a result, common scoring matrices within aligners like GraphAligner and minimap 2 (Li 2021) prefer the more complete, but wrong, haplotype (Supplementary Figure S2), leaving unresolved gaps in the assembly. To address this, we use a two fold strategy of (1) improvements to GraphAligner and (2) supplemental alignments from Winnowmap (Jain *et al*. 2022).

First, GraphAligner was modified to use a new diploid heuristic for read alignment. This heuristic is based on finding homologous pairs of nodes and the heterozygous *k*-mers which distinguish them, matching the read to a specific haplotype based on matching heterozygous *k*-mers and preventing the read from aligning to nodes in the other haplotype. Despite the name, the diploid heuristic does not specifically use haplotype information and instead deals with alignment in repeats with copy count two in general, which most commonly are the two haplotypes of a diploid genome. This section uses the terms *heterozygous* to refer to any unique, one-copy sequence in the genome and *homozygous* for any two-copy sequence.

To detect heterozygous *k*-mers, first all closed syncmers (Edgar 2021) are found and counted from the node sequences with parameters *k* and *s*, and are used as the *k*-mers for the rest of the diploid heuristic. Any heterozygous *k*-mer must have a copy count of exactly one. However, a *k*-mer with copy count of one may not necessarily be a heterozygous *k*-mer. This includes cases where one of the haplotypes has a gap due to low LA coverage or assembly issues, in which case any homozygous *k*-mers in the remaining haplotype will have a copy count of one in the graph.

Homozygous, two-copy *k*-mers are used to identify homology between different nodes. After the *k*-mer copy counts are tabulated, GraphAligner detects substrings which are enclosed by two-copy *k*-mers on both ends, and contain only one-copy *k*-mers in between. If two such substrings are found in different nodes such that the enclosing two-copy *k*-mers match, the substrings are considered to be a homologous pair. Since the substrings are enclosed by *k*-mers with copy count of two, any such substring can match with at most one other substring, but since the nodes are of variable length, multiple short nodes can be homologous to a single long node. All homologous pairs are detected in the graph and stored in an index. In addition, the heterozygous *k*-mers inside the homologous substrings, as well as their locations, are stored as a list of haplotype informative *k*-mers.

When aligning a read, the diploid heuristic is used to decide which nodes should be used based on their *k*-mer matches. Haplotype informative *k*-mers are found in the read and clustered based on their match diagonal between the read and the nodes. If at least three haplotype informative *k*-mers from the same node are found within a 100 *b* diagonal, the node is considered to match the read at the corresponding location. All other nodes that are homologous to the matched node are also checked. If there is even one haplotype informative *k*-mer match to an another homologous node, then the pair is considered ambiguous and ignored. If there are no haplotype informative *k*-mer matches to the other node (i.e. it appears to be from a different ‘haplotype’), the read is forbidden from aligning to the unmatched node in the region covered by the matching node.

In the seed-and-extend phase of alignment, when a read is forbidden from aligning to a node, any seed hits found between the read and the forbidden node are discarded, and the dynamic programming algorithm is not allowed to extend the alignment into a forbidden node. Importantly, the alignment is only forbidden within the region covered by the matching node, since inexact, two-copy repeats within the same haplotype often have homologous substrings, and forbidding the alignment over the entire read would cause issues in such regions.

The diploid algorithm requires syncmer parameters *k* and *s*. Higher values of *k* will find more haplotype informative *k*-mers in repetitive areas and requires higher accuracy reads in order to find matches, while a lower *k* will find fewer haplotype informative *k*-mers but enables the heuristic to work with higher error reads. GraphAligner allows using multiple *k* values which will run the diploid heuristic separately for each *k* and then merge the lists of forbidden nodes. Parameter *s* is always set to five and *k* defaults to *k* = 21 and *k* = 31 for the two rounds of the diploid heuristic. Future higher accuracy UL reads may benefit from higher values of *k*.

Lastly, Winnowmap alignments are used as a supplement to the GraphAligner alignments, since Winnowmap was designed for haplotype-specific alignments (but is not graph aware). In this phase, all tips in the graph are identified along with their alternate haplotype nodes. Any reads aligned by GraphAligner to these nodes are then extracted and realigned to the full graph using Winnowmap. The resulting read alignments are used for gap filling in cases where the Winnowmap alignments differ from GraphAligner and connect two otherwise disconnected tip nodes. Reads not used for gap filling in this step have their alignments reverted to the original GraphAligner alignments to allow for their use in other graph simplification operations.

### Hi-C data processing

Verkko2 uses Hi-C chromosome conformation capture data for both phasing and scaffolding of the assembly graph, and supports both short-read Hi-C as well as Nanopore-based Pore-C (Deshpande *et al*. 2022) sequencing data. Critically, Verkko integrates this long-range linkage information directly with the assembly graph, improving accuracy over standalone phasing and scaffolding tools.

The two main features that make Hi-C data useful for T2T assembly are that (1) Hi-C read pairs are more frequently derived from the same haplotype than between different haplotypes, and (2) Hi-C read pairs are more frequently derived from close rather than distant positions in the genome. These features are based on the 3D organization of the genome and make Hi-C data particularly useful for phasing (Selvaraj *et al*. 2013) and scaffolding (Burton *et al*. 2013), respectively.

Figure 2 summarizes the new steps added to Verkko pipeline to enable Hi-C-based phasing and scaffolding of Verkko’s ULA assembly graph. Hi-C read pairs are first aligned to the assembled unitigs with BWA-MEM (Li 2013). All read pairs where at least one read is mapped ambiguously (mapping quality of 0, typically multi-mappers from homozygous regions of different haplotypes) are ignored for phasing but used later in scaffolding. For the alignment of the long-read Pore-C data, we use Minimap2 (Li 2021) and consider each pair of fragments in a read separately. Otherwise, the handling of Pore-C data mirrors that of Hi-C.

**Fig. 2:**
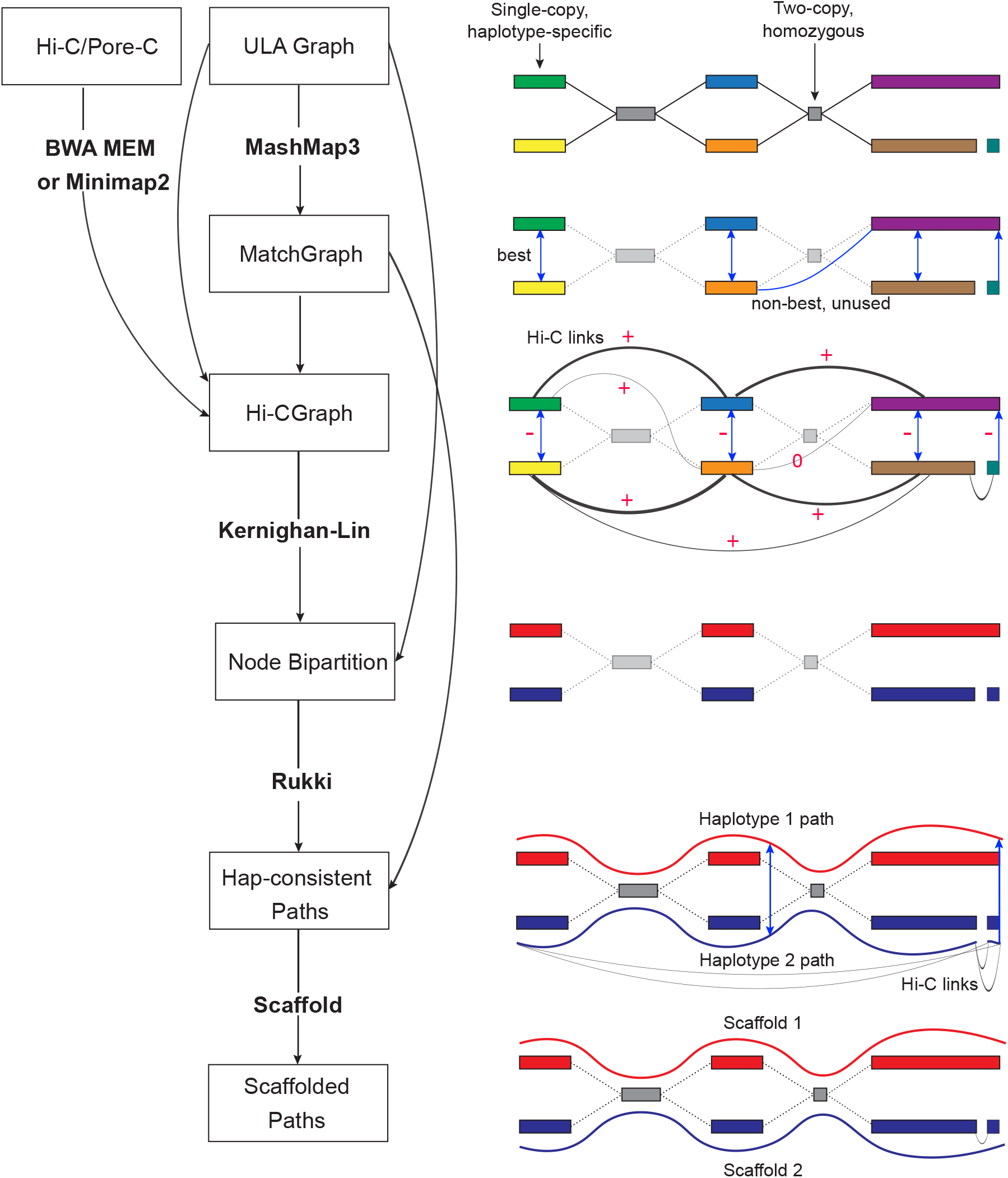
Overview of the Verkko2 Hi-C/Pore-C processing. The process starts with the ULA graph built from the LA and UL sequences. Note that the ULA graph links are only used to cluster the graph into connected components so they are shown in gray in the figure. The BWA/Minimap2 step aligns the Hi-C or Pore-C data to the sequences of the ULA graph nodes and counts connecting pairs of nodes are tallied. Next, the Match Graph step ignores homozygous (based on coverage) and short (by default ≤ 200 *kb*) nodes. The remaining nodes are self-aligned to identify homology. The initially computed Hi-C edges are filtered using to build the Hi-C Graph. In many cases, the highest count Hi-C connection is between homologous pairs of nodes. To avoid these false links, edges connecting potentially repetitive nodes are removed (shown with a value of 0) while edges connecting homologous nodes are given large negative weights (shown with −). These updated link weights are used to bi-partition each connected component of the graph into two haplotypes. These partitions are then provided to Rukki along with the ULA graph to generate haplotype paths and the pipeline proceeds as in Verkko1. These haplotype-consistent paths are again used to identify homology, shown with blue edges based on the MatchGraph step. All four possible connections are considered to connect the two blue paths based on Hi-C link evidence. In this example, the alignment of both blue paths to the red path adds a multiplicative bonus to one Hi-C connection consistent with the alignment, leading to a scaffold connecting the blue paths.

### Hi-C haplotype phasing

Due to the fact that Hi-C read pairs more often originate from the same halpotype, the frequency of Hi-C links between assembly graph nodes can be used to phase (i.e. partition) the graph by haplotype. However, because of Hi-C library noise, sequencing errors, and mapping issues, homologous nodes from different haplotypes can also share a high number of Hi-C links. If only considering the frequency of Hi-C links between nodes, this can pose a problem for phasing.

To resolve the issue of Hi-C links between homologous nodes, we construct an additional set of edges on top of the ULA graph that are based on sequence similarity between the nodes, called the MatchGraph. Homologous nodes in the ULA graph are identified by an all-vs-all similarity search using MashMap3 (Kille *et al*. 2023), and all pairs of nodes with at least one homology match over a minimum length threshold (default 200 *kb*) are connected by an edge in the MatchGraph. The weight of each edge is set to the total bases covered by alignments between the corresponding nodes.

One might expect that a node would be connected only to the corresponding homologous node of similar size. However, this is not the case in real assembly graphs, since some regions may be assembled in one node in one haplotype but split into multiple nodes in the other. Worse, some relatively long regions between heterologous chromosomes can share sequences that are similar in identity to that of homologous chromosomes, e.g. centromeres, rDNAs, pseudoautosomal regions (PAR), and pseudo-homologous regions (PHRs) (Nurk *et al*. 2022; Guarracino *et al*. 2023). To address this issue, the edges of the MatchGraph are processed in two different ways to distinguish homology between *haplotypes* and *repeats*. For each node *V* in the MatchGraph, we check whether there is an edge incident to *V* with the weight being a clear majority (default *>* 1.5 * *second best weight*) among all edges incident to *V* and also greater than one third of the length of at least one of those nodes. Such edges are considered to link a pair of haplotypes, while edges of the MatchGraph that are not best for either of their incident nodes are considered to link a pair of repeats.

Hi-C read pair alignments are then considered along with the MatchGraph to build the HiCGraph. The HiCGraph is constructed from long nodes (default *>* 200 *kb*) of the ULA graph, with undirected, weighted edges corresponding to the number of Hi-C read pairs mapping between a pair of nodes. To correct for erroneous Hi-C links between homolgous nodes, nodes connected by a haplotype edge in the MatchGraph are given a large negative weight (default −10 * *maximumedgeweight*) in the HiCGraph, and to correct for mismapped Hi-C pairs between repeats, nodes connected by a repeat edge in the MatchGraph are removed from the HiCGraph. Additionally, self-edges are removed since they carry no additional phasing information.

The resulting HiCGraph is then phased by identifying a bipartition of the nodes that minimizes the total weight of edges between two similarly-sized subsets. Although this problem is known to be NP-complete (Garey 1997), the Kernighan-Lin algorithm (Kernighan and Lin 1970) is a commonly used greedy heuristic implemented by the NetworkX Python library (Hagberg *et al*. 2008). It operates by iteratively swapping pairs of nodes that yield the largest decrease in edge weight between the two partitions. We begin with multiple (default 1, 000) randomly generated bipartitions and perform 10, 000 iterations of the Kernighan-Lin algorithm for each component. Afterwards, any short nodes not included in the initial HiCGraph are assigned based on whether a clear majority (*best* ≥ 2.5\**second_best*) of Hi-C links connect it to one of the two subgraphs.

Although it is possible to phase the full HiCGraph at once, we split it into subgraphs that are expected to represent diploid chromosome components prior running the Kernighan-Lin algorithm and process each individually. These components are defined by supplementing the ULA assembly graph with the haplotype edges of the MatchGraph, after which any Hi-C links connected nodes from different components are ignored. This helps to reduce the number of iterations of Kernighan-Lin and also enables the detection and proper handling of heterogametic sex chromosomes. After phasing, the bipartitions of each component are arbitrarily merged to generate a single bipartition of the entire ULA graph, which is then passed to the Rukki module to identify haplotype paths in the same manner as a trio-based bipartition (Rautiainen *et al*. 2023). Supplementary Figure S3 summarizes all steps of the Hi-C phasing process in detail.

### Haplotype path reconstruction

Rukki is an independent Verkko module for extracting haplotype paths from a labeled assembly graph. Given an assembly graph and node labels corresponding to haplotype information (either trio or Hi-C based), Rukki classifies the nodes into maternal, paternal, or homozygous categories and then performs a heuristic search for haplotype paths starting from long nodes in the graph. The homozygous nodes are assumed to be sequences of perfect similarity between the two haplotypes and are included in both. The sequences of the resulting haplotype paths can be extracted from the graph, forming contigs (gapless) and scaffolds (gapped).

In Verkko2, Rukki has been improved to minimize the number of nodes that remain unassigned to a haplotype. First, the criteria for initial assignment of haplotypes is relaxed to ensure that all large nodes (default *>* 500 kb) with appropriate coverage are assigned. Second, all nodes in a tangle, where the tangle is connected to only a single haplotype, are assigned to the same haplotype. Third, for ambiguous bubbles where the haplotype separation is unclear, Rukki now forces the two paths of the bubble into different haplotypes using any available phasing information, rather than including a single path in both haplotypes as before (which resulted in a loss of heterozygosity). Lastly, Rukki now provides a text description for the causes of scaffold gaps, including: graph tangle, coverage dropout, or ambiguous bubble with inadequate phasing information.

### Hi-C scaffolding

In addition to using Hi-C for phasing, Verkko2 introduces the use of Hi-C data for scaffolding to connect haplotype paths that remain broken due to missing edges in the assembly graph (e.g. due to low sequencing coverage) or separated in the graph by very large tangles (e.g. due to the rDNA arrays). Scaffolding is performed directly on the haplotype-specific paths identified by Rukki in the preceeding step.

Hi-C interaction strengths between each pair of haplotype paths are computed based on Hi-C read pairs mapped within a fixed-length prefix/suffix of the paths (default 5 Mb or half the path length, whichever is smaller) and scores are computed for all for possible orientations of the pair (fwd-fwd, fwd-rev, rev-rev, rev-fwd). To account for multi-mapping reads, we filter out read pairs that have at least one read with more than five mappings of equivalent quality, or both reads in the pair have at least one mapping to the same graph node.

The scaffolding step considers two types of Hi-C link weights, a *primary* weight using all read pairs including the remaining multi-mappers normalized by their number of alignments (i.e. if one read in pair is mapped to two places and second to three, each link is counted with weight 1/6), and a *secondary* weight using only uniquely mapped reads. Additional scaffolding information contained within the assembly graph is incorporated via multiplicative bonuses on the link weights. First, paths that are close (closer than 500 kb) in the assembly graph are more likely to be consecutive and their corresponding link weights are multiplied by a constant factor (default three). Second, it is common for a sequencing gap to be present in only one of the two homologous chromosomes, so, using the previously constructed MatchGraph, we check if paths *p*_1_ and *p*_2_ are homologous haplotypes of some third path *p*_3_ and whether the position of these matches are close in *p*_3_. If so, a constant multiplicative bonus (default five) is applied. This homology bonus can also be applied using alignments to a reference genome, as a form of reference-assisted scaffolding.

Given the Hi-C link weights as computed above, scaffolds are constructed via the connection of mutually best pairs of paths. For simplicity, both orientations of a haplotype path are considered independently but forbidden from appearing in the same scaffold. For each path *p*_1_ that does not end with a telomere, we check for the existence of a clear best forward extension path *p*_2_ and also require that *p*_1_ is a clear backward extension for *p*_2_ (*best >* 1.5 * *second best*). If there is no mutual extension using the primary weights, we check the secondary weights using the same rule.

Lastly, subtelomeric repeat complexity and sequencing dropout can lead to scaffolds that are missing a small telomere-containing node at their ends. To resolve such cases, we perform telomere-informed graph scaffolding in cases where one haplotype is T2T but the homologous haplotype appears broken. Given an assembly graph component where one Hi-C path *p*_1_ contains telomere on both ends and the homologous path *p*_2_ contains telomere only on one end, if (1) there is a path *p*_*t*_ that also contains telomere on one end, (2) *p*_*t*_ is near (closer than 500 kb) *p*_2_ in the graph, and (3) no other paths containing nodes from this component longer than 1 Mb, then *p*_2_ and *p*_*t*_ are scaffolded together.

### Assembly post-processing

The output of the scaffolding module is an updated set of paths through the ULA graph, which are converted to sequences through the existing Verkko consensus procedure (Rautiainen *et al*. 2023). Verkko2 adds the option to generate a BAM file describing the positions and alignments of the underlying reads. These alignments are informed by knowledge contained within the assembly pipeline, such as where LA read *k*-mers were used to construct nodes and where UL reads were aligned to the graph, resulting in a more accurate representation than if the reads were naively mapped to the resulting assembly. This output is useful for both inspecting the quality of the resulting consensus sequence and for potentially further improving the quality of the consensus through raw read, signal-level polishing (Mastoras *et al*. 2024b).

To facilitate easier submission to the public sequence archives, Verkko2 also includes support for the removal of small, unassembled sequences, as well as the isolation of specific, user-defined sequences. For human assemblies, default screening sequences are provided for mitochondrial, rDNA, and Epstein-Barr virus (EBV) sequences (NC 012920 (Andrews *et al*. 1999), KY962518 (Kim *et al*. 2018), and AJ507799 (Arrand *et al*. 1981), respectively), but there are no restrictions on what sequences can be used and this feature can also be used for the removal of any known contaminant.

To screen the final assembly, first, nodes with no connection to any other node in the graph are identified and removed if shorter than a minimum length (default *<* 100 kb). Second, all sequences in the assembly are compared against the screening sequences using MashMap3. Mapping results are filtered to remove those with an estimated match identity below 97.5% or those that cover less than 10% of the sequence. If the remaining hits, combined, cover at least half of the sequence(s), it is declared a match and removed from the assembly. Additionally, any node within four hops of a match is also declared a match if its sequence is not more than 4− fold longer than the matched node sequence, by default. This captures more diverged versions of the screened sequence and excludes incomplete resolutions of repetitive arrays unless they are included in a large node. For all matched contigs, an exemplar sequence is reported as the sequence with the largest breadth of coverage by matches, or the match with the highest LA read coverage when multiple contigs exist with matches covering more than 90% of their length. Finally, since circular molecules (e.g. mitochondria) are often assembled into linear contigs with redundant sequence on their ends (Hunt *et al*. 2015), the extracted exemplar is trimmed to remove any redundancy *>* 1 kb.

## Results

### Benchmarking

We selected two human datasets from the Human Pangenome Reference Consortium (HPRC, Liao *et al*. (2023)) for detailed benchmarking. We selected datasets with both Hi-C and parental trio data available for comparison and to allow trio-based validation. Although newer data is available for HG002, we used the same data that was used in Rautiainen *et al*. (2023) to highlight the algorithmic improvements rather than sequencing technology evolution. To measure Verkko’s performance on species other than human we included a sheep cross (of *O. canadensis* and *O. aries*) from the Ruminant T2T Consortium (Kalbfleisch *et al*. 2024; Olagunju *et al*. 2024) and a chicken from the Vertebrate Genomes Project (Rhie *et al*. 2021). Supplementary Table S1 summarizes the read and genome information for all included datasets.

We report the number of T2T contigs and scaffolds for measuring contiguity; hamming and switch error rates for phasing correctness estimated with yak (https://github.com/lh3/yak); Phred-scaled quality value (QV) for consensus accuracy measured with yak; and compleasm (Huang and Li 2023) for completeness at the gene level with the haplotypes evaluated independently, corrected for sex chromosome genes, and values averaged between the haplotypes. T2T contigs and scaffolds are both defined by the presence of telomeric sequence on the ends, but with scaffolds allowing for gaps (typically caused by large tandem repeat arrays). Comparison of Verkko2’s trio-based and Hi-C-based phasing are shown in Table 1, demonstrating comparable accuracy between the two methods. To avoid overcounting the number of missing genes for heterogametic samples, missing genes were also counted for the autosomal sequences as identified by MashMap3 alignment to the existing reference genomes (CHM13 for human (Nurk *et al*. 2022), GCA 024206055.1 (Huang *et al*. 2023) for chicken, and GCF 016772045.2 (Davenport *et al*. 2022) for sheep).

**Table 1.** Verkko2 Hi-C vs trio benchmarking. T2T contigs/scaffolds were counted as those longer than 5 Mb with telomeres on both ends. Contigs were broken into scaffolds at any stretch of *>* 3 Ns, and contigs/scaffolds shorter than 100 kb were discarded for all metrics. Verkko Hi-C shows comparable QV, switch, hamming, and missing gene stats to Verkko trio, but it consistently has a higher count of T2T scaffolds due to its ability to restore missing connectivity using Hi-C links.

|  | sheep | chicken | HG002 | HG00733 |
| --- | --- | --- | --- | --- |
| T2T scf Hi-C | **31** | **34** | **40** | **41** |
| T2T scf trio | 23 | 32 | 32 | 33 |
| T2T ctg Hi-C | **24** | 21 | 21 | **26** |
| T2T ctg trio | 20 | **25** | **22** | 23 |
| Hamming err Hi-C | **0.85%** | **0.58%** | **0.39%** | **0.75%** |
| Hamming error trio | 0.88% | 0.60% | **0.39%** | 0.77% |
| Switch err Hi-C | **0.95%** | **0.13%** | **0.41%** | **0.79%** |
| Switch error trio | **0.95%** | **0.13%** | **0.41%** | **0.79%** |
| QV Hi-C | **54.17** | 45.13 | 53.87 | **53.86** |
| QV trio | **54.17** | **45.17** | **53.89** | 53.82 |
| missing genes Hi-C | 1.37% | 3.12% | 1.61% | **0.09%** |
| missing genes trio | **1.36%** | **2.52%** | **1.60%** | **0.09%** |
| missing genes (no sex chrs) Hi-C | **0.06%** | 1.06% | **0.09%** | **0.09%** |
| missing genes (no sex chrs) trio | **0.06%** | **0.44%** | **0.09%** | **0.09%** |
| dup genes Hi-C | 1.73% | **0.12%** | **0.68%** | **0.71%** |
| dup genes trio | **1.72%** | 0.22% | 0.69% | 0.72% |

We next compared Verkko2 against Verkko1 and Hifiasm (Cheng *et al*. 2021), which was recently updated to support hybrid assembly with ultra-long reads (Cheng *et al*. (2024), version 0.19.8). Figure 3 summarizes key metrics with the full results provided in Supplementary Table S2. Verkko2 shows clear improvements in T2T scaffolds over both tools, with similar correctness results, and a dramatically reduced runtime compared to Verkko1. Verkko2 is now also faster than Hifiasm, but with runtime dependent on the sequencing coverage and repeat structure of each genome. Hamming error rate results are similar with the exception of Hifiasm Hi-C which is less accurate than other tools. Focusing on the distal regions of the human acrocentric chromosomes, Verkko2 Hi-C resolves five and six out of the ten possible acrocentric chromosomes for HG002 and HG00733, respectively, while no other assemblies resolve more than one. In the sheep *O. aries* haplotype, where the majority of chromosomes are telocentric and share sheep satellite I repeats at one end (D’Aiuto *et al*. 1997), Verkko1 resolved only four chromosomes, including Chromosomes 3 and X, which are known to be deficient in this satellite (Burkin *et al*. 1996). In contrast, Verkko2 Hi-C resolved an additional six T2T scaffolds for this challenging *O. aries* haplotype.

**Fig. 3:**
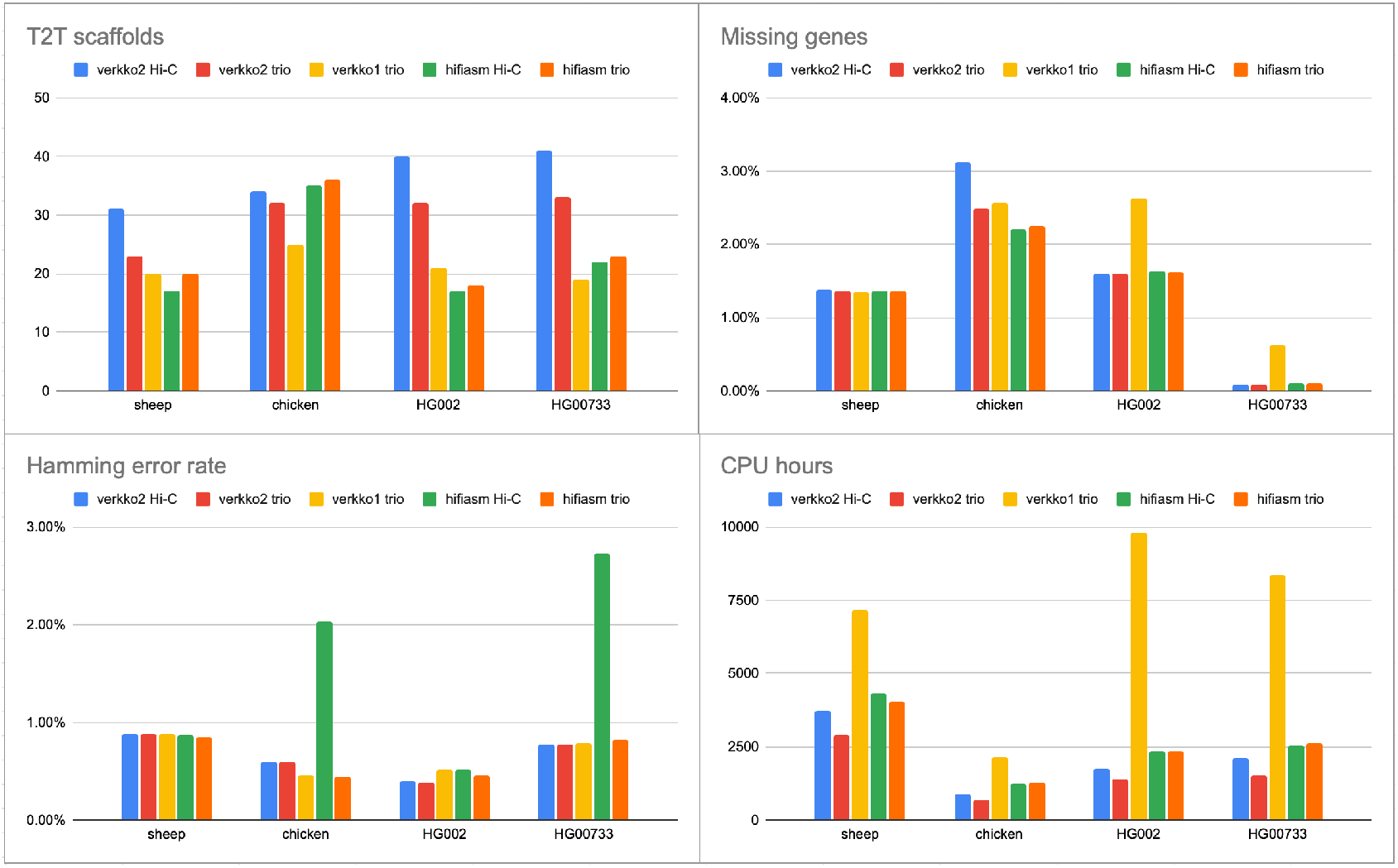
Comparison of tested assemblers with all statistics measured as before. Verkko Hi-C has the highest T2T scaffold rate (except on chicken), followed by Verkko2 trio. Both versions of Hifiasm are comparable to Verkko1. With the exception of Hifiasm Hi-C on chicken and HG00733, all assemblers have comparable hamming error rates. Verkko1 has a consistently higher rate of missing genes compared to other assemblies. All assemblers were run on the NIH Biowulf compute cluster.

The above reference-free metrics demonstrate Verkko2’s improvements for both human and non-human genomes, but none of these metrics are able to capture large, structural errors in the assemblies. Existing reference-free tools such as Flagger (Liao *et al*. 2023) and NucFrec (Vollger *et al*. 2019) produce a high number of false-positive predictions and are not always consistent with each other (Porubsky *et al*. 2024), making it challenging to use them without manual curation of their results. A high-quality T2T reference genome exists for HG002 but, while reference-based validation (Gurevich *et al*. 2013) is commonly used to evaluate the quality of haploid genomes, there is not yet a standard tool for evaluating the quality of large, diploid assemblies. We use QUAST (Gurevich *et al*. 2013; Mikheenko *et al*. 2018) which, while not specifically designed for diploid assemblies, is suitable to perform a relative comparison of base-level and structural correctness. For all metrics reported, Verkko2 trio is the most accurate with Verkko2 Hi-C a close second (Supplementary Table S3). Verkko2 also reduces misassemblies by almost twofold versus Verkko1 and has approximately 2.5-fold fewer base level errors than Hifiasm.

### HPRC Year 1 reassembly

To evaluate Verkko2’s performance across a larger set of samples, we selected a subset of 17 HPRC samples (Liao *et al*. 2023) for which both trio and Hi-C data was available (Figure 4, Supplementary Table S4). These Verkko2 HPRC assemblies have a median T2T scaffold count of 39, median hamming and switch error rates below 0.6%, and a median QV of 54.94. The core gene statistics can identify both missing or mis-assigned sequences. For example, if homologus sequences are placed into the same haplotype, it would have an increase in duplicated genes while the other haplotype would have an increase in missing genes. If only one haplotype has an increase in missing genes, the sequence for it likely incomplete but was not placed in the other haplotype. However, the assemblies are stable with a median missing gene rate of 0.17%. One sample (HG02559) was assembled into 45 T2T scaffolds, one short of the ideal 46 for a diploid human genome, while 40% of all samples have more than 40 T2T scaffolds. We confirmed the absence of chimeric assemblies by aligning all Verkko scaffolds ≥ 500 kb to the T2T-CHM13 reference genome (Nurk *et al*. 2022) using MashMap3 and found no scaffold with *>* 1% of its sequence mapped to more than one chromosome. (excluding the chrX/Y PAR and acrocentric chromosomes, see below). In comparison, the published HPRCv1 assemblies from (Liao *et al*. 2023), which did not incorporate the UL data, achieved only two T2T scaffolds (compared to 39 for Verkko2); median hamming and switch error rates of 0.7% and 0.6%; median missing genes per haplotype of 0.23%; and a median QV of 53.6 (Liao *et al*. 2023). Thus, Verkko2’s incorporation of UL and Hi-C data not only improves T2T assembly continuity but also reduces overall assembly errors.

**Fig. 4:**
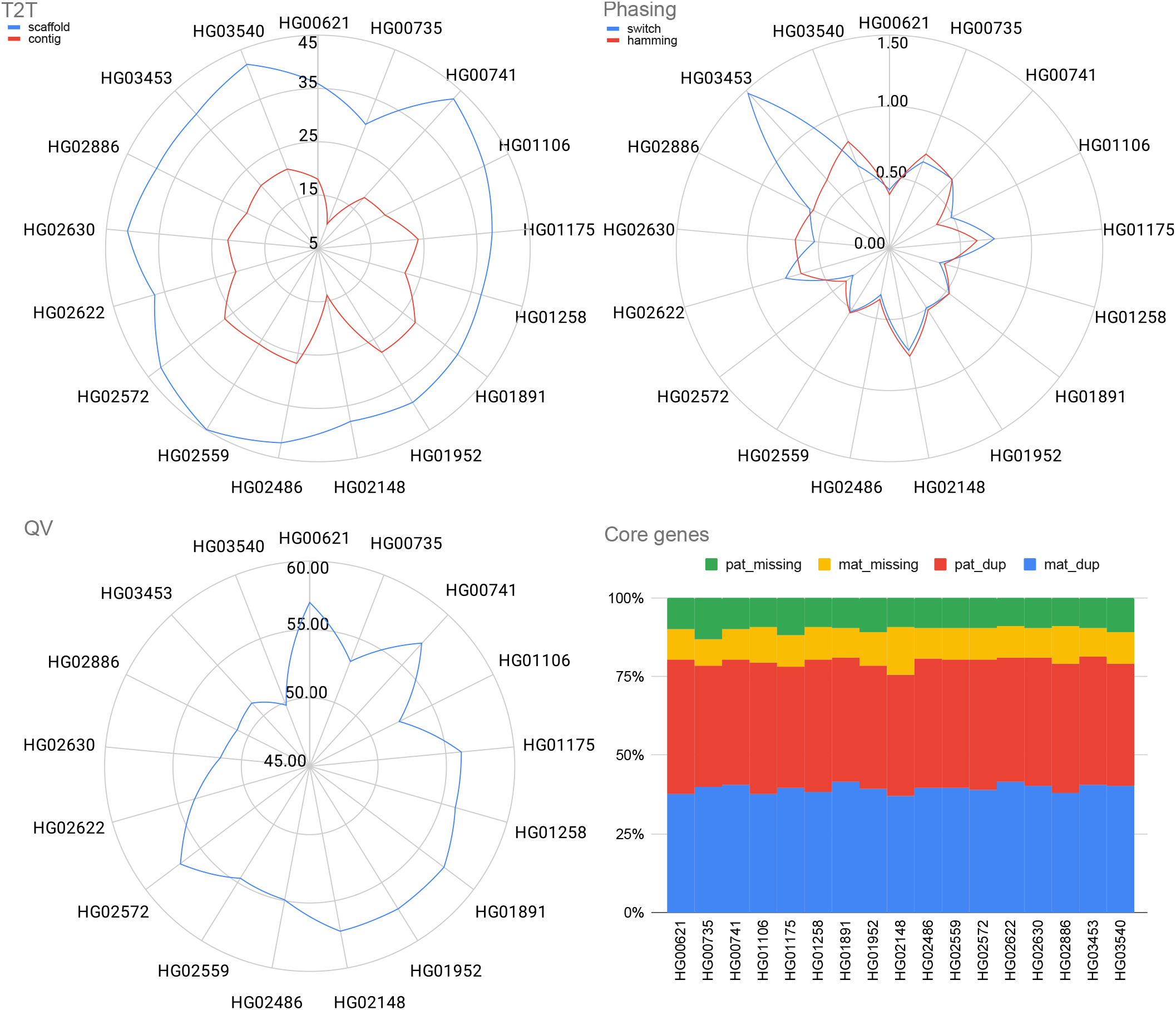
Verkko2 results on the HPRC year 1 assemblies. The T2T statistics are computed as before. QV and phasing error are computed using yak and the average of both haplotypes is reported. The core genes are computed using compleasm and reported for each haplotype. The dup categories report single-copy genes present more than once in a haplotype. Due to natural variation, a small number (*<* 1%) of duplicated genes is expected. The missing gene categories report single-copy genes not present in the assembly, excluding the sex chromosomes. The stability of duplicated and missing genes across all samples supports that Verkko2 is accurately reconstructing the full sequence for both haplotypes.

This large set of human trio samples also provided the opportunity to further measure our Hi-C scaffolding accuracy, specifically within the difficult-to-validate short arms of the acrocentric chromosomes. These short arms share regions of sequence similarity above 99% identity between heterologous chromosomes (Nurk *et al*. 2022) and undergo meiotic recombination (Guarracino *et al*. 2023), confounding sequence alignment and ruling out reference-based validation. Additionally, these chromosomes are known to co-localize within nucleoli, potentially violating our assumptions of Hi-C interaction patterns. To validate Verkko2’s scaffolding performance within these difficult regions, we relied on the trio information to identify haplotype switches in the acrocentric scaffolds. While this evaluation will not detect all scaffolding errors (e.g. *maternal* Chromosome 13 short arm swapped with *maternal* Chromosome 14 short arm), we would expect a random scaffold join within the acrocentrics to introduce a haplotype switch with approximately 50% probability (e.g. *maternal* Chromosome 13 short arm swapped with *paternal* Chromosome 13 short arm). Using Merqury (Rhie *et al*. 2020), we detected the parental origin (maternal/paternal/unassigned) for each long node (*>* 1 Mb) and quantified haplotype switches between the distal region of the short arm and the rest of the acrocentric chromosome (spanning the rDNA array). In total, Verkko2 scaffolded 93 acrocentric chromosomes and introduced no haplotype switches between the distal region and the rest of the chromosome. A single switch, occurring within the distal region, was observed on a short arm of sample HG03453. Thus, the absence of haplotype switches observed across the rDNA array within these 17 HPRC samples suggests that our Hi-C scaffolding method is generally accurate. For the HG002 genome, where distal regions have been experimentally validated with fluorescence in situ hybridization (Potapova *et al*. 2024b), Verkko2 scaffolded five distal regions, which were all consistent with the reference.

## Discussion

The Verkko2 improvements described here dramatically improve the number and quality of T2T chromosomes assembled, while simultaneously reducing computational requirements. Combined, these changes approach the goal of complete, diploid T2T assembly (e.g. 46 T2T chromosomes per human genome, see Supplemental Table S4). For the most difficult to assemble human acrocentric chromosomes, Verkko2 was able to resolve 55% of all chromosomes tested as T2T scaffolds with only the rDNA array typically remaining as a gap. Assembling across these most repetitive regions of the genome will enable future research on their unique recombination patterns (Guarracino *et al*. 2023), and will improve our understanding of the evolutionary history and structural variation within these important chromosomes across large pangenome datasets such as the HPRC (Liao *et al*. 2023).

To simplify sequencing requirements, Verkko2 now supports multiple long-read datatypes, in addition to HiFi, for initial construction of the LA multiplex De Bruijn graph. This includes ONT Duplex (Koren *et al*. 2024; Sarashetti *et al*. 2024) reads as well as ONT Simplex reads that have been sufficiently corrected (Stanojevic *et al*. 2024). Verkko2 now also provides base-level alignments and location information for all reads used to generate the assembly. This can be combined with existing validation and polishing tools (Liao *et al*. 2023; Mastoras *et al*. 2024a; Mc Cartney *et al*. 2022) to incorporate the assembler’s representation of the genome and is a first step towards routinely reporting quality values for all bases in the assembly.

Areas for future improvement include better utilization of depth of coverage information, improved assemblies in the absence of UL data, pseudo-haplotype outputs in the absence of phasing data, and support for polyploid genomes. Although Verkko2 is more tolerant to coverage fluctuations, coverage is still not used in the most optimal way. Verkko2 only uses coverage from uniquely placed LA nodes, leading to systematic underestimates in repetitive regions. Improved coverage estimates, specifically the identification of unique nodes and the estimated copy number of repeat nodes, would improve overall assembly continuity. Second, Verkko’s pipeline is currently optimized for the hybrid combination of LA and UL reads. In cases where only LA data is available, graph simplification steps currently only performed on the ULA graph, could be performed directly on the LA graph, improving results in the absence of UL data. Third, Verkko2 only outputs completely phased contigs and thus produces more fragmented assemblies when no long-range information (such as trio, Hi-C, or Strand-seq (Henglin *et al*. 2024)) is available. In this case, a pseudo-haplotype graph traversal that allows for phase switches, while preserving the overall structure of the assembly, would improve continuity. Lastly, Verkko2’s scaffolding could be further improved by considering multiple suffix/prefix lengths rather than a single fixed value, as is done in YAHS (Zhou *et al*. 2023); considering multi-way interactions from Pore-C libraries rather than only pairwise; and extending the phasing/scaffolding methods to polyploid genomes.

The routine assembly of complete genomes promises to improve all aspects of genomics. For example, the current paradigm for human clinical genomics involves mapping reads against a single reference genome and identifying the variants between the sample and reference. However, this approach is fundamentally flawed due to reference bias and limits our understanding of the genome, especially in the context of multi-copy gene families and other repetitive regions. Inferring the complete, diploid genome of an individual, at a cost that is comparable to current mapping-based approaches, promises to correct this deficiency (Ebler *et al*. 2022). Until then, Verkko2’s input flexibility, fast runtime, and novel scaffolding methods enable the routine assembly of personalized reference genomes and the construction of a comprehensive T2T pangenome database. This will expand the fraction of the genome and types of complex variation that are captured by large-scale genomic studies and surveyed by clinical diagnostic tests, bringing us one step closer to a future of personalized, telomere-to-telomere genomics.

## Author contributions statement

DA, MR, SN, BPW, SS, SK developed and implemented algorithmic improvements, DA developed the new scaffolding module and performed benchmarking. MR developed the new overlapper module. SK supervised software development. AMP directed the project. DA, SK, and AMP drafted the manuscript with methods contributions from MR, SN, and SS. All authors read and approved the final manuscript.

## Dataset availability

No new data was generated for this study. Sheep reads were generated by ruminant T2T consortium (Kalbfleisch *et al*. 2024)

Reads for chicken sample were downloaded from https://www.genomeark.org/genomeark-all/Gallus_gallus.html for individual bGalGal1 and its parents bGalGal2 and bGalGal3. All reads for human samples were downloaded from Human Pangenome Reference Consortium’s Amazon cloud, https://s3-us-west-2.amazonaws.com/human-pangenomics/index.html

## Acknowledgments

The authors would like to thank the HPRC assembly working group and Shilpa Garg for helpful discussions on Hi-C alignment and phasing methods. We would like to thank the Ruminant T2T consortium for sharing the bighorn sequencing data and Wesley Warren (USDA NIFA 2022-67015-36218), Erich Jarvis, and the Vertebrate Genome Project for sharing the chicken data. This work was supported, in part, by the Intramural Research Program of the National Human Genome Research Institute, National Institutes of Health (DA, BPW, SS, AMP, and SK). This work utilized the computational resources of the NIH HPC Biowulf cluster (https://hpc.nih.gov) and CSC - IT Center for Science Finland.

