## Supplementary Materials for "Verkko2: Integrating proximity ligation data with long-read De Bruijn graphs for efficient telomere-to-telomere genome assembly, phasing, and scaffolding"

### 1. TABLES

Table S1
Table S2
Table S3
Table S4

| | sheep | chicken | HG002 | HG00733 |
|---|---|---|---|---|
| Genome size (Gb) | 2.7 | 1.1 | 3.1 | 3.1 |
| Het Rate (%) | 0.988 | 0.950 | 0.262 | 0.114 |
| HiFi | | | | |
| N50 | 23,393 | 21,912 | 13,607 | 15,537 |
| Total Bases (Gb) | 213.33 | 109.35 | 198.30 | 183.05 |
| ONT | | | | |
| Bases in reads >=100 kb (Gb) | 105.09 | 17.75 | 95.60 | 170.84 |
| Total Bases (Gb) | 497.32 | 177.00 | 247.19 | 271.14 |
| Hi-C | | | | |
| Total Bases (Gb) | 65.19 | 116.46 | 125.87 | 198.95 |

**Table S1.** Information about datasets used for benchmarking. Heterozygosity level was estimated with genomescope [1] using the HiFi reads, except for chicken, where genomescope crashed on HiFi data. For that sample Hi-C Illumina reads were used for estimation. Heterozygosity of the heterogametic samples (sheep, chicken, HG002, HG01109) can be overestimated with this tool.

| Species | T2T scf | T2T ctgs | Hamming error | Switch error | QV | Missing genes | Missing genes (no sex chr) | CPU Hours | Peak Memory |
|---|---|---|---|---|---|---|---|---|---|
| **Sheep** | | | | | | | | | |
| Verkko2 Hi-C | **31** | **24** | **0.85**% | **0.58**% | 54.17 | 1.37% | **0.06%** | 3725.80 | 206 |
| Verkko2 trio | 23 | 20 | **0.85**% | 0.95% | 54.17 | **1.36**% | **0.06%** | **2897.02** | 203 |
| Hifiasm Hi-C | 17 | 16 | 0.86% | 0.95% | 57.25 | 1.37% | **0.06%** | 4342.36 | 381 |
| Hifiasm trio | 20 | 19 | **0.85%** | 0.94% | **57.46** | **1.36**% | **0.06%** | 4046.67 | 396 |
| Verkko1 trio | 20 | 15 | **0.85**% | 0.95% | 55.85 | **1.36%** | **0.06%** | 7181.07 | **196** |
| **Chicken** | | | | | | | | | |
| Verkko2 Hi-C | 34 | 21 | 0.58% | **0.13%** | 45.13 | 3.12% | 1.07% | 870.31 | 84 |
| Verkko2 trio | 32 | 25 | 0.58% | **0.13%** | **45.17** | 2.52% | 0.47% | **673.58** | 85 |
| Hifiasm Hi-C | 35 | 32 | 2.01% | 0.33% | 40.34 | **2.20%** | **0.15%** | 1252.55 | 202 |
| Hifiasm trio | **36** | 35 | **0.41%** | 0.34% | 40.25 | 2.25% | 0.20% | 1268.15 | 203 |
| Verkko1 trio | 25 | 23 | 0.43% | 0.30% | 39.88 | 2.56% | 0.49% | 2150.19 | **69** |
| **HG002** | | | | | | | | | |
| Verkko2 Hi-C | **40** | 21 | 0.39% | **0.41%** | 53.87 | 1.61% | **0.09%** | 1736.25 | **164** |
| Verkko2 trio | 32 | **22** | **0.38%** | **0.41%** | 53.89 | **1.60%** | **0.09%** | **1394.57** | **164** |
| Hifiasm Hi-C | 17 | 9 | 0.51% | 0.47% | 55.12 | 1.64% | 0.13% | 2347.53 | 325 |
| Hifiasm trio | 18 | 10 | 0.46% | 0.48% | **55.29** | 1.61% | 0.11% | 2330.15 | 328 |
| Verkko1 trio | 21 | 8 | 0.46% | 0.50% | 51.52 | 2.64% | 1.13% | 9794.19 | 165 |
| **HG00733** | | | | | | | | | |
| Verkko2 Hi-C | **41** | **26** | 0.75% | **0.79%** | 53.86 | **0.09%** | **0.09%** | 2112.23 | 165 |
| Verkko2 trio | 33 | 23 | **0.74%** | **0.79%** | 53.82 | **0.09%** | **0.09%** | **1518.56** | 165 |
| Hifiasm Hi-C | 22 | 14 | 2.73% | 0.86% | **56.63** | 0.10% | **0.09%** | 2552.40 | 283 |
| Hifiasm trio | 23 | 15 | 0.81% | 0.87% | 56.52 | 0.10% | 0.10% | 2629.20 | 275 |
| Verkko1 trio | 19 | 11 | 0.78% | 0.83% | 51.97 | 0.62% | 0.61% | 8345.69 | **162** |

**Table S2.** Comparison of tested assemblers on human and non-human data on all metrics. Scaffolds $<$ 100 kb were discarded for all metrics. T2T scaffolds are scaffolds longer than 5 Mb that contain telomeres (detected by seqtk telo) on both ends. Hamming error rate, switch error rate, and QV were calculated with yak. Missing genes count were calculated with compleasm v0.2.6 (haplotypes evaluated independently, average values reported). All assemblers were run on the NIH Biowulf cluster. Best values for each metrics and sample are highlighted in bold. Verkko2 Hi-C has the highest T2T scaffold count with the exception of chicken where it is two less than the best. Verkko2 trio has the lowest runtime across all datasets, followed by Verkko2 Hi-C. While Verkko1 has the lowest memory usage, Verkko2 only modestly increases memory while reducing runtime $2.5 - 7$-fold.

| | Verkko2 Hi-C | Verkko2 trio | Verkko1 trio | Hifiasm Hi-C | Hifiasm trio |
|---|---|---|---|---|---|
| NA50 | 133.576 | **133.990** | 130.098 | 95.008 | 101.268 |
| NA90 | 45.180 | **45.332** | 36.924 | 39.271 | 39.310 |
| misassemblies | 97 | **63** | 119 | 277 | 163 |
| Genome fraction % | **99.91** | **99.91** | 99.7 | 98.60 | 99.78 |
| local misassemblies | 216 | **214** | 550 | 816 | 420 |
| mismatches per 100Kbp | 0.43 | **0.41** | 1.33 | 2.96 | 1.06 |
| indels per 100Kbp | 0.77 | **0.75** | 0.92 | 1.28 | 0.87 |
| N's per 100 kbp | 34.15 | **18.21** | 55.43 | 185.77 | 81.77 |

**Table S3.** QUAST accuracy evaluation of all tested assemblies on HG002 dataset, using HG002 genome release v1.1 as a reference. Scaffolds $<$ 100 kb were discarded for all metrics. NA50 and NA90 are reported in Mb. Best values among all assemblers is highlighted in bold.

| Sample ID | T2T ctg | T2T scf | Hamming | Switch | QV | Missing | Missing (no sex chr) |
|---|---|---|---|---|---|---|---|
| **HG00621** | 18 | 36 | 0.41% | 0.38% | 57.00 | 1.72% | 0.17% |
| **HG00735** | 10 | 30 | 0.65% | 0.71% | 53.26 | 0.21% | 0.20% |
| **HG00741** | 18 | 43 | 0.66% | 0.65% | 57.19 | 0.17% | 0.17% |
| **HG01106** | 19 | 40 | 0.48% | 0.37% | 52.29 | 1.70% | 0.17% |
| **HG01175** | 24 | 38 | 0.74% | 0.62% | 56.17 | 0.20% | 0.20% |
| **HG01258** | 22 | 37 | 0.36% | 0.40% | 56.06 | 1.70% | 0.17% |
| **HG01891** | 28 | 38 | 0.53% | 0.52% | 57.28 | 0.16% | 0.16% |
| **HG01952** | 28 | 39 | 0.49% | 0.51% | 57.24 | 1.72% | 0.19% |
| **HG02148** | 14 | 38 | 0.73% | 0.77% | 57.23 | 0.25% | 0.25% |
| **HG02486** | 27 | 42 | 0.33% | 0.36% | 54.94 | 1.69% | 0.16% |
| **HG02559** | 26 | 45 | 0.52% | 0.53% | 54.67 | 0.16% | 0.16% |
| **HG02572** | 27 | 42 | 0.32% | 0.38% | 56.80 | 1.69% | 0.16% |
| **HG02622** | 21 | 37 | 0.76% | 0.65% | 53.74 | 0.16% | 0.16% |
| **HG02630** | 22 | 41 | 0.53% | 0.66% | 51.56 | 0.16% | 0.16% |
| **HG02886** | 20 | 39 | 0.62% | 0.60% | 50.94 | 0.19% | 0.19% |
| **HG03453** | 21 | 39 | 1.48% | 0.65% | 51.29 | 0.16% | 0.16% |
| **HG03540** | 21 | 42 | 0.62% | 0.80% | 49.82 | 0.18% | 0.18% |
| Verkko2 Hi-C Median | 21 | 39 | 0.53% | 0.60% | 54.94 | 0.20% | 0.17% |
| Hifiasm yr1 Median | 0 | 0 | 0.71% | 0.61% | 53.57 | 0.24% | 0.23% |

**Table S4.** HPRC Yr1 assembly metrics for Verkko2 Hi-C. T2T scaffolds are scaffolds longer than 5 Mb that contain telomeres (detected by seqtk telo) on both ends. Hamming error rate, switch error rate, and QV were calculated with yak. Missing genes count were calculated with compleasm v0.2.6 (haplotypes evaluated independently, average values reported).

#### 2. FIGURES

Figure S1
Figure S2
Figure S3

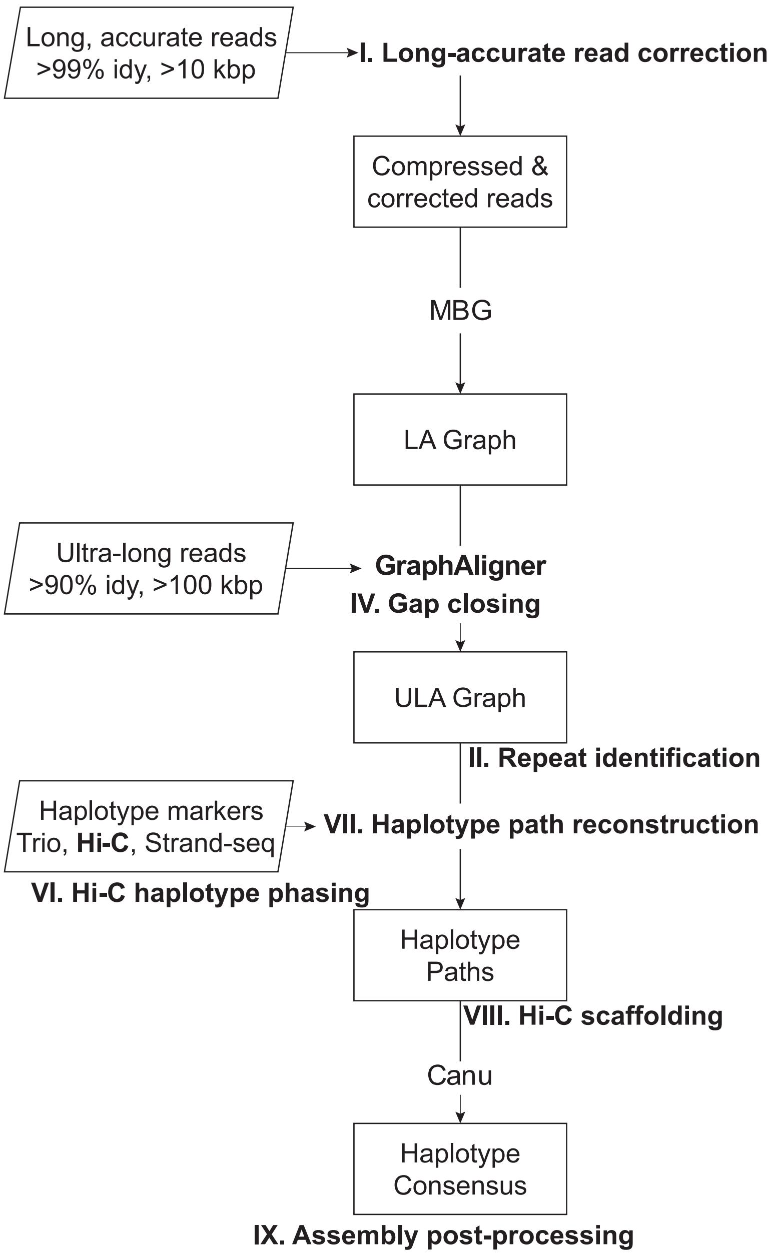


**Fig. S1.** Verkko1 pipeline graphical representation, adapted from [2]. Stages modified in Verkko2 are labeled with roman numerals and are described in the corresponding subsections of Methods: I: Long-accurate read correction, II+III: Repeat identification and better assembly for telomeres, IV: Gap closing, VI: Hi-C haplotype phasing, VII: Haplotype path reconstruction, VIII: Hi-C scaffolding, and IX: Assembly post-processing.

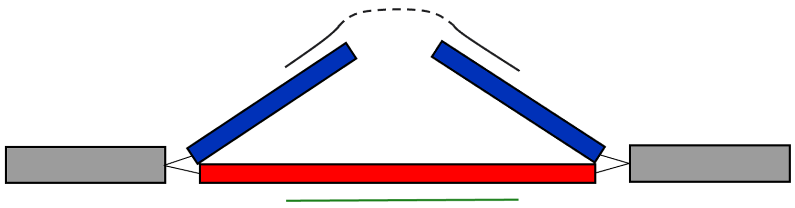

**Fig. S2.** An example of two possible alignments of an ONT read to a gapped region of the LA graph. The grapy nodes are homozygous and used by both haplotypes. The red and blue nodes correspond to the maternally-inherited and paternally-inherited haplotype, respectively. The paternally-inherited haplotype has a gap due to a coverage dropout in the LA data. The correct alignment is represented by two solid black lines connected by a dash. The middle part of the read sequences comes from a region absent in the LA graph and is represented by dashed black line. The alignment to the alternate haplotype is represented by solid green line. This haplotype does not have missing sequence in the LA graph. Although the alternate haplotype may have lower identity, it can have a higher score and be selected because it provides a single alignment with more bases covered by the alignment and no gap penalties.

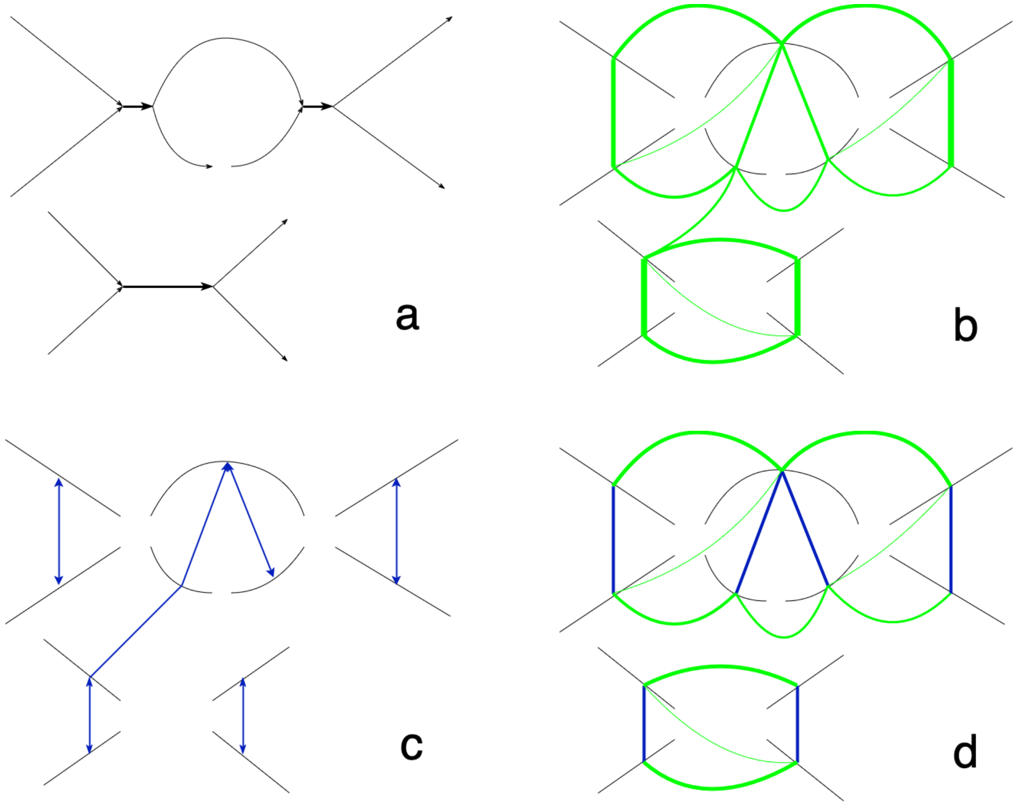


**Fig. S3.** Steps of Hi-C the phasing algorithm: a) Initial ULA assembly graph with two connected components. b) The Hi-C Graph prior to any filtering. The thickness of edges correspond to the number of Hi-C read pairs mapping to both nodes c) The MatchGraph, with alignment matches shown in blue. The arrow on the edges indicates best matches. For example an arrow pointing from node $x$ to node $y$ to show that $y$ is the best match for $x$. d) The filtered Hi-C Graph. The MatchGraph edges are used to generate large negative weights (shown in blue). Non-best edges (e.g. connecting the two components) are set to have a 0 value and are dropped in this figure. Remaining Hi-C edges are shown in green.
